# ActiveDriverDB: human disease mutations and genome variation in post-translational modification sites of proteins

**DOI:** 10.1101/178392

**Authors:** Michal Krassowski, Marta Paczkowska, Kim Cullion, Tina Huang, Irakli Dzneladze, B. F. Francis Ouellette, Joseph T. Yamada, Amelie Fradet-Turcotte, Jüri Reimand

## Abstract

Interpretation of genetic variation is required for understanding genotype-phenotype associations, mechanisms of inherited disease, and drivers of cancer. Millions of single nucleotide variants (SNVs) in human genomes are known and thousands are associated with disease. An estimated 20% of disease-associated missense SNVs are located in protein sites of post-translational modifications (PTMs), chemical modifications of amino acids that extend protein function. ActiveDriverDB is a comprehensive human proteo-genomics database that annotates disease mutations and population variants using PTMs. We integrated >385,000 published PTM sites with ∼3.8 million missense SNVs from The Cancer Genome Atlas (TCGA), the ClinVar database of disease genes, and inter-individual variation from human genome sequencing projects. The database includes interaction networks of proteins, upstream enzymes such as kinases, and drugs targeting these enzymes. We also predicted network-rewiring impact of mutations by analyzing gains and losses of kinase-bound sequence motifs. ActiveDriverDB provides detailed visualization, filtering, browsing and searching options for studying PTM-associated SNVs. Users can upload mutation datasets interactively and use our application programming interface for pipelines. Integrative analysis of SNVs and PTMs helps decipher molecular mechanisms of phenotypes and disease, as exemplified by case studies of disease genes *TP53*, *BRCA2* and *VHL*. The open-source database is available at https://www.ActiveDriverDB.org.

## Introduction

Genome-wide association and DNA sequencing studies have enabled large-scale characterisation of human genomes and revealed millions of single nucleotide variants (SNVs), copy number alterations, and other types of genetic variants. Interpreting these vast datasets and identifying genotype-phenotype associations, molecular mechanisms, causal disease variants and cancer driver mutations remain major challenges of current biomedical research (1,2). To date, tens of thousands of genomes of healthy individuals and cancer samples have been analysed with DNA sequencing to catalogue protein-coding variation in large-scale genomics projects such as The Cancer Genome Atlas (TCGA)(3), the Interactional Cancer Genome Consortium (ICGC) (4), the 1,000 Genomes Project (5), The Exome Aggregation Consortium (ExAC) (6), and others. Further, comprehensive open-access databases such as ClinVar (7) collect clinically annotated disease genes and mutations. The functional impact of SNVs is studied using a family of scoring and machine learning methods such as PolyPhen2 (8), SIFT (9) and CADD (10) that distinguish benign and deleterious variants by analysing evolutionary sequence conservation and other genomic features. These tools and resources are used to prioritize candidate SNVs. However information about protein signalling and interactions is not routinely applied to understand the genetic basis of human variation and disease.

Post-translational modifications (PTM) are chemical modifications of amino acids that act as molecular switches and expand the functional repertoire of proteins. Post-translational regulation of proteins is carried out by modular reader-writer-eraser networks where specific enzymes induce PTMs in target proteins, remove PTMs, and interact with modified sites (11). There are more than 400 known types of PTMs that can be mapped systematically with high-throughput proteomics technologies (12,13). Four types of PTMs have the currently largest experimental datasets available: phosphorylation, ubiquitination, acetylation and methylation, with nearly 400,000 sites available in public databases (14-16). PTMs are involved in various aspects of cellular organization, including protein-protein interactions, protein activation and degradation, regulation of chromatin state, organismal development, and signaling pathways associated with human disease and cancer (17-20). Further, PTMs are increasingly targetable by drugs and used in precision cancer therapies (21-23). Thus PTM information helps interpret genetic variation, map genotype-phenotype associations, and investigate molecular mechanisms of disease.

PTM sites are enriched in known disease mutations, cancer driver mutations, and rare population variants (24-29). Sequence motifs in PTM sites recognized by enzymes such as kinases are also frequently mutated in disease (25,30), leading to losses of existing sequence motifs and acquisition of new motifs, with potential rewiring of cellular signaling networks (27,31). However, common methods for predicting functional impact of SNVs such as PolyPhen2 (8), SIFT (9) and CADD (10) do not directly account for PTMs and often underestimate the impact of variants in PTM sites, including known disease mutations (25). The PhosphositePlus database maintains downloadable datasets with PTM site variation (14), however a dedicated comprehensive database of genetic variation in PTM sites does not exist to our knowledge.

To address this limitation, we developed the ActiveDriverDB database, a proteo-genomics resource for interpreting human genome variation using information on post-translational modification sites in proteins (24-27). The database integrates experimental evidence about PTMs from recent studies of somatic cancer mutations from large-scale genomics studies (3)[submitted: *Automating Somatic Mutation calling for Ten Thousand Tumor Exomes. Ellrott K., Covington K. R., Kandoth C., Saksena G., McLellan M, Bailey M.H., Sofia H., Hutter, C., the MC3 Working Group and the TCGA Network*], mutations in known disease genes (7), and human genome variation from population studies (5,32). We also display the network context of PTM SNVs by analysing protein-protein interactions specific to PTM sites as well as drugs targeting upstream PTM enzymes (33). The database allows browsing, visualising and interpreting hundreds of thousands of genome variants predicted to affect PTM sites in proteins. Besides comprehensive annotation and visualization of SNVs derived from large genomics datasets, users can interactively upload, store and analyze their own custom datasets of mutations. The database is designed for interactive study of genes and mutations and hypothesis generation for basic and translational biologists, while computational biologists can also take advantage of downloadable datasets and access the database automatically through an application program interface (API). Our database is open source and can be downloaded for local use.

### Genomic and proteomic data in ActiveDriverDB

For interpreting genome variation using protein PTMs, ActiveDriverDB includes two major types of human biomolecular data: genomics data on missense SNVs and proteomics data on PTMs (**Table 1**). The terminology of genes and proteins is used interchangeably in the manuscript, however in most cases we refer to the primary protein isoform of the gene defined by the HGNC gene symbol.

**Table 1.** Overview of genome variation datasets and post-translational modifications included in ActiveDriverDB. Counts of PTMs and missense SNVs reflect all high-confidence protein isoforms collected in the database.

|  |  | TCGA<br>PanCanAtlas | ClinVar | 1000<br>Genomes | ESP6500 | Total |
| --- | --- | --- | --- | --- | --- | --- |
| **Dataset** |  |  |  |  |  |  |
|  | Size | >11,000 exomes | 494,059 records | 2504 genomes | 6503 exomes | - |
|  | Description | Cancer (somatic) | Inherited disease | General population |  | - |
| **Mutations** |  |  |  |  |  |  |
|  | Total | 1,841,901 | 138,938 | 1,071,308 | 1,319,079 | 3,818,672 |
|  | in PTM sites | 257,333 | 27,116 | 143,986 | 185,920 | 540,676 |
|  | with network-rewiring effect | 36,393 | 3,357 | 20,704 | 26,678 | 75,598 |
|  | Annotated nucleotides (hg19/GRCh37) | 2,158,851 | 155,824 | 1,211,581 | 1,486,067 | 4,401,677 |
| **PTM sites affected by mutations\*** |  |  |  |  |  |  |
|  | Total | 233,716 | 16,487 | 157,772 | 186,736 | 309,407 |
|  | Phosphorylation sites | 184,864 | 12,914 | 128,372 | 150,451 | 244,034 |
|  | Acetylation sites | 12,684 | 1,251 | 7,236 | 8,984 | 16,779 |
|  | Ubiquitination sites | 38,325 | 2,641 | 22,738 | 28,220 | 51,320 |
|  | Methylation sites | 3,564 | 272 | 2,298 | 2,577 | 4,499 |
| **Proteins with mutations affecting PTM sites** |  |  |  |  |  |  |
|  | Total | 27,861 | 3,317 | 25,149 | 26,202 | 29,542 |
|  | Kinases & PTM enzymes | 548 | 101 | 516 | 529 | 556 |
|  | Kinase families | 127 | 59 | 127 | 126 | 127 |
| **PTM sites** |  |  |  |  |  |  |
|  | Total | - | - | - | - | 385,185 |
|  | Phosphorylation sites | - | - | - | - | 299,241 |
|  | Acetylation sites | - | - | - | - | 21,670 |
|  | Ubiquitination sites | - | - | - | - | 67,933 |
|  | Methylation sites | - | - | - | - | 5,666 |

Human genome variation datasets include disease-associated SNVs and those attributed to the human population in general. First, ActiveDriverDB includes somatic cancer mutations of nearly 10,000 tumor samples of 34 types derived from large-scale exome sequencing projects conducted by the TCGA. Somatic cancer mutations are derived from the recent MC3 release of the TCGA PanCanAtlas project [submitted: *Ellrott K., et al.*]. Known and putative inherited disease mutations from the ClinVar database (7) are also available in ActiveDriverDB. Missense SNVs representing inter-individual human genome variation from large-scale population sequencing projects are also integrated into the database, including the 1,000 Genomes Project (5) with more than 2,500 genomes and the ESP6500 project (32) with more than 6,500 exomes.

Proteomics data on post-translational modification sites in human proteins is based on published studies and is retrieved from public databases PhosphositePlus(14), Phospho.ELM(15) and HPRD(16). The database uses four types of PTMs with the largest number experimentally determined sites available – 299,241 phosphorylation sites, 67,933 ubiquitination sites, 21,670 acetylation sites and 5,666 methylation sites with 385,185 distinct sites in total counted across all protein isoforms. Site-specific protein-protein interactions of PTM enzymes and their substrate proteins are also derived from these databases and comprise experimentally predicted kinase-substrate interactions as only few such interactions are known for other enzymes and modification types. We also integrate drugs that target upstream PTM enzymes from the DrugBank database (33) to provide translational hypotheses to researchers interpreting PTM-associated SNVs.

In total, the database characterises 540,676 unique missense SNVs in PTM sites across high-confidence protein isoforms, including 257,333 in cancer genomes, 27,116 in inherited disease and more than a hundred thousand variants in population sequencing projects. Above a quarter of unique SNVs in high-confidence cancer genes from the Cancer Gene Census database(34) are associated with PTM sites (26% or 29,243 SNVs). Among disease genes annotated in the ClinVar database, 27,071 unique missense SNVs (19%) are associated with PTM sites. These statistics suggest that a large fraction of germline and somatic disease mutations can be interpreted using PTM information.

### Estimating the impact of mutations on PTM sites

To discover SNVs in PTM sites, we implemented the analysis pipeline used in our previous studies (24-27). In brief, genomic coordinates of mutations were mapped to substitutions in protein sequences using the Annovar software and RefSeq genes of human genome assembly version hg19/GRCh37. Peptide sequences corresponding to PTM sites were derived from public databases (14-16) and mapped to corresponding RefSeq protein sequences using exact sequence matching and by permitting multiple matches per sequence. PTM sites extended seven amino acids upstream and downstream of the modified protein residue, and multiple clustered PTM sites were merged into consecutive regions. Protein domains were retrieved from the InterPro database (35) and mapped into non-redundant regions. Disorder predictions were derived using the DISOPRED2 software (36). ActiveDriverDB provides information for nearly 30,000 high-confidence isoforms of human genes, however primary isoforms according to the Uniprot database (37) are shown by default.

Missense SNVs (i.e., substitutions) in PTM sites are annotated with further information regarding their position relative to PTMs. PTM SNVs are considered direct if they substitute the central amino acid residue that is subject to post-translational modification. Indirect PTM SNVs are classified either as proximal or distal (1-2 or 3-7 amino acid residues to the nearest PTM site, respectively). We also distinguish variants that affect the four types of PTM sites (phosphorylation, acetylation, ubiquitination, methylation) and mutations affecting multiple adjacent PTM sites of different types.

We also perform sequence motif analysis with our machine learning method MIMP (27) to predict impact of mutations on kinase binding sites. MIMP analyses mutations in PTM sites using 476 models of sequence motifs bound by 322 kinases and families of kinases derived from public databases and experimental datasets (14-16,38,39). It predicts whether SNVs in the motifs cause loss of existing kinase-bound motifs or creation of *de novo* kinase-bound motifs. These predictions suggest the impact of missense SNVs on the rewiring of cellular signalling networks.

### Searching and browsing PTM mutations in proteins

The database provides a flexible graphical user interface for finding, visualizing and interpreting SNVs in PTM sites and their potential impact on signaling networks.

#### Searching for genes, pathways, and diseases

The main search bar available on the front page and on the top of each page is a flexible multi-search bar that supports several search objectives. First, the user can identify a gene (protein) of interest by either its HGNC symbol that retrieves the primary isoform by default (*e.g.*, *TP53*) or a RefSeq transcript ID that retrieves a specific protein isoform (*e.g., NM_000546*). Second, the database supports search for biological processes of Gene Ontology(40), molecular pathways of Reactome (41) (*e.g., Wnt signaling pathway*). and diseases in the ClinVar database (*e.g., Noonan syndrome*). The latter options retrieve lists of genes ranked by the number of PTM SNVs. These search options benefit researchers who are interested in interpreting specific genes, pathways, or disease mutations.

#### Searching for mutations

The user can search for a gene (protein) according to a mutation of interest. This option supports both protein amino acid substitutions (*e.g., TP53 R282W*) as well as missense variants shown in chromosomal coordinates (*e.g., chr17 7577094 C T*). The latter option is supported by our rapid indexing system that covers all potential missense SNVs in the human genome. Search for mutations is especially beneficial for genetics researchers who have identified interesting missense SNVs in a genome-wide association or DNA sequencing study.

#### Browsing top disease-associated genes and pathways

In addition to focused searches, users may browse sets of disease-associated genes and pathways with unexpectedly frequent mutations in PTM sites, indicating potential molecular mechanisms of disease. Candidate gene lists are available for cancer mutations from TCGA and inherited disease mutations from ClinVar. Disease gene lists are ranked according to statistical enrichment of mutations in PTM sites estimated using our ActiveDriver method (26). These lists are useful for novice users of the database who are looking for an overview of the database through examples of genes and pathways.

### Visualisation and analysis of PTM SNVs

The two primary views in the database, *the protein sequence view* and *the interaction network view*, are focused on individual genes (proteins). Both views provide detailed visualizations of PTM-associated SNVs, tables with additional information, protein descriptions, and external links. The views permit filtering of mutations by dataset (inherited disease mutations, somatic cancer mutations, or variants in population genomics studies), disease types, and PTM types. All non-PTM mutations can be filtered as well. Both views display the primary isoform as default, while alternative isoforms can be selected.

### Protein sequence view: sequence features, mutation hotspots, SNV impact on PTMs and network rewiring

The protein sequence view comprises three main components: the needleplot of mutation frequencies and predicted impact PTM sites, integrated sequence tracks with protein domains and disorder, and a detailed table of mutations with frequencies and predicted impact. The needleplot shows the linear view of the protein sequence of interest and provides a detailed overview of SNVs in the protein and their impact on post-translational modifications (**Figure 2A**). Mutations are shown as needles extending vertically from the sequence and PTMs are shown as blocks of blue tones on the sequence. High-frequency mutations are shown with taller needles, and SNVs with PTM impact are colored. Mouse-over motion over needle heads displays information about each individual mutation, its disease annotations, known PTM enzymes such as kinases binding the site, predictions of network rewiring with mutation-induced gains and losses of sequence motifs (**Figure 2B**), as well as drugs targeting the upstream kinases and other PTM enzymes. The needleplot can be zoomed and searched by amino acid position. Mutations are also shown below the needleplot as a table that can be sorted and filtered (**Figure 2C**). The protein view additionally displays integrated sequence tracks with the protein amino acid sequence, protein domains from the InterPro database (35), and protein sequence disorder predictions from the DISOPRED2 software (36). The needleplot can be exported as a high-resolution image in PDF format and the mutation table can be exported as a text file or a spreadsheet.

**Figure 1.**
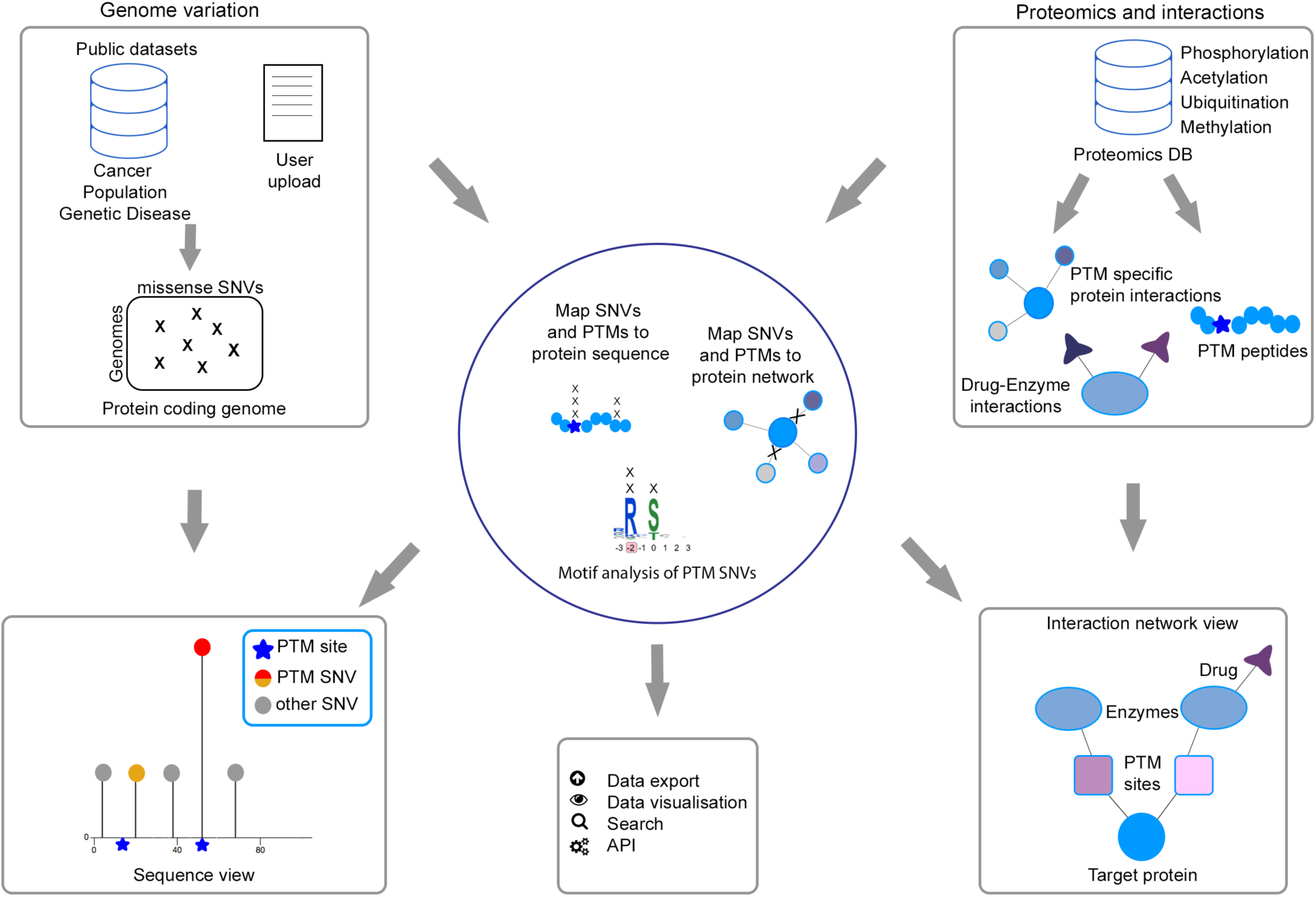
Overview and workflow of ActiveDriverDB. Our proteo-genomic database resource integrates genomic and proteomic data for interpreting human genome variation and disease mutations with information on post-translational modifications (PTMs). Genomics datasets (top-left panel) include cancer genome sequencing studies (TCGA), disease genes and mutations (ClinVar), and human genome variation studies (ESP6500, the 1,000 Genomes Project). Proteomics datasets (top-right panel) include PTM sites of four commonly studied PTM types derived from public databases (PhosphositePlus, Phospho.ELM, HPRD), site-specific interactions of PTM enzymes and target proteins, and drug interactions with PTM enzymes. Our systematic analysis pipeline (middle panel) aligns PTM sites with missense single nucleotide variants (SNVs), predicts the impact of SNVs on kinase-bound sequence motifs using the MIMP software, and organizes SNVs, PTMs and upstream enzymes into site-specific interaction networks. The protein sequence view of the database (bottom left panel) shows the distribution of PTMs and SNVs along the protein sequence, while interaction network view (bottom right panel) shows the site-specific interactions of mutated proteins with upstream PTM enzymes and their associated drugs. The database also provides various exporting, visualization, searching and automated analysis tools (bottom middle panel).

**Figure 2.**
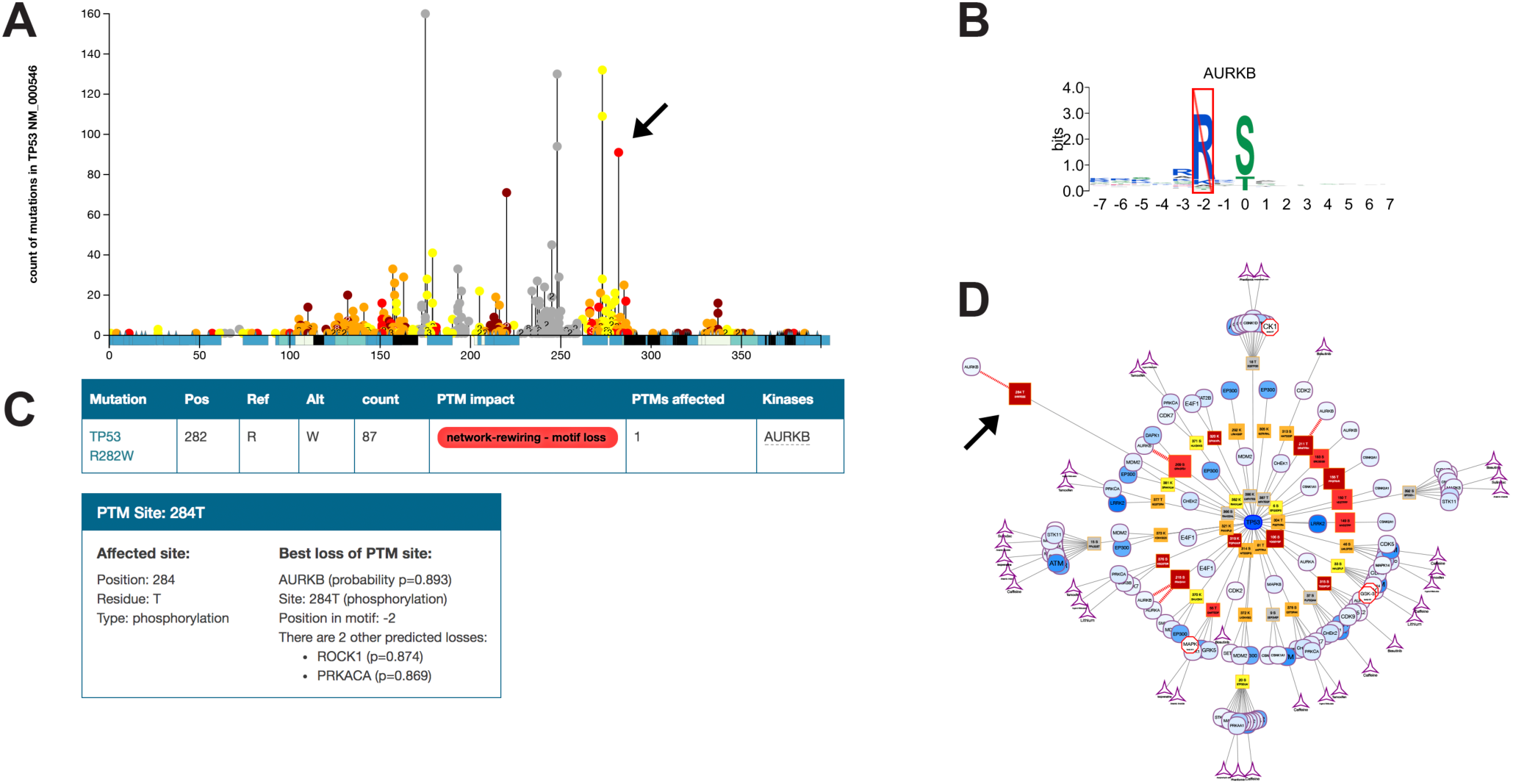
PTM-associated cancer mutations in the tumor suppressor protein TP53. **(A)** Needleplot in the protein sequence view shows the distribution of missense cancer mutations from TCGA (vertical bars) and their associations with PTM sites (blue boxes) with protein sequence on the x-axis and number of mutations on the Y-axis. **(B)** The substitution R282W disrupts the sequence motif bound by the Aurora Kinase B (AURKB). **(C)** Table of mutations with detailed information, disease associations, and impact on SNVs. **(D)** The experimentally determined interaction network shows the TP53 protein (middle node) and its PTM site-specific interactions with upstream PTM enzymes, as well as approved drugs targeting these enzymes. Node shapes indicate types of interacting molecules and sites: protein of interest (oval), PTM sites (square), enzymes interacting with PTM site (circle), and drugs targeting the enzyme (triangle). Arrows indicate the interaction of TP53 and Aurora kinase B at phosphosite T284 and the associated substitution R282W.

### Interaction network view: protein interactions in PTM sites, upstream enzymes and drug targets

In the interaction network view, the selected protein is displayed with the interactions of enzymes known to bind its PTM sites. These sub-networks show the protein of interest, all its enzyme-bound PTM sites, and the enzymes binding each site (**Figure 2D**). Drugs targeting the upstream enzymes are also shown. Upstream enzymes and drug targets are useful for generating translational hypotheses for follow-up experiments such as drug screens in mutated cell lines. The majority of interactions comprise phosphorylation sites and associated kinases, as most site-specific interactions are known for this PTM type. To highlight interesting proteins in the interaction network, upstream enzymes are colored according to their mutation frequency. The interaction networks are visualized using an automatic layout algorithm that emphasizes the hierarchical structure of the network. The network layout is interactive and can be zoomed and manually rearranged for clarity. The network can be exported as a text file and as a high-resolution PDF image.

Two interaction networks are available for each protein. The high-confidence experimental network comprises site-specific kinase-substrate interactions that have been determined in experimental studies. The computationally predicted network is based on sequence motif analysis of experimentally derived phosphorylation sites using the MIMP software. The latter network shows kinase-substrate interactions that are gained and lost through site-specific mutations in motifs as predicted by the MIMP software.

### Interactive analysis of custom datasets of mutations

Users can upload their own datasets of genome variants as VCF or tab-separated files. This option allows users to interpret their unpublished and published datasets using PTM sites and to directly compare their findings with public datasets. The uploaded variants are stored for short term in a password-protected area of the website and most visualization and database features are available for analysis. The database supports upload of DNA variants as chromosomal coordinates and amino acid substitutions as protein-level coordinates.

### Automated database access using the application progamming interface (API)

ActiveDriverDB includes a simple application programming interface (API) based on the Representational State Transfer (REST) pattern. This state-of-the-art interface allows computational biologists and data scientists access the data through multiple programming environments like R and Python. The queries include protein or DNA-level coordinates of mutations and the ActiveDriverDB returns PTM annotations of input variants, after converting DNA coordinates to protein coordinates.

Additional options for queries are available, including filters for mutation types (cancer, inherited disease, population), querying of mutations by gene symbol or refSeq ID, and others described in documentation online.

### Software design and availability

The web application uses the Flask Micro framework and two relational databases: the first for constant biological data and the second for dynamic content and user-provided data. Additionally a key-value BerkleyDB database is used for mapping of all potential missense SNVs in the human genome. Visualisations are implemented in the d3.js framework. Our needleplots are inspired by the original muts-needle-plot library (https://zenodo.org/record/14561). All code is available on terms of LGPL 2.1 license. Detailed technical documentation is available on GitHub at https://github.com/reimandlab/ActiveDriverDB.

### Frequent cancer mutations in PTM sites in the tumor suppressor protein TP53

The tumor suppressor *TP53* is mutated in 50% of all cancers. The transcription factor TP53 regulates the gene expression of numerous cellular processes by binding specifically to DNA. Consistently, most cancer mutations in TP53 are located in the DNA-binding domain (DBD) of the protein and one third of these mutations are clustered in seven hotspot regions (R175, G245, R248, R249, R273, R282) (**Figure 2A**) (42). Although most of the mutations in TP53 lead to inhibition of its transcriptional activity and loss of function, mutations such as R282W lead to gain of function and provide TP53 with oncogenic abilities that are distinct from its wild-type roles (reviewed in (42,43)). To date, the mechanisms by which R282W leads to this transformation are still unclear (44).

Interestingly, MIMP analysis of sequence motifs in TP53 predicts that the substitution of R282 to tryptophan, glycine or glutamine induces a network reviewing effect by abolishing the sequence motif of the Aurora B kinase (AURKB) in the phosphosite T284 of TP53 (**Figure 2B, 2C**) (27). *In vitro* phosphorylation assays as well as immunoprecipitation experiments of TP53 in cells with ectopic expression of Aurora B show that the kinase interacts and phosphorylates TP53 on multiple residues, including T284 (**Figure 2D**) (27,45,46). Consistent with a role of the kinase in regulating TP53 activity, substitution of T284 for a glutamic acid inhibits it ability to promote the transcription of *CDKN1A* (45). According to our analyses, more than 250 mutations in the TCGA dataset have the potential to affect phosphorylation at T284 in a direct, proximal or distal manner (**Table 2**), suggesting that these mutations may regulate a common cancer-related function of TP53. As R282W is associated with early cancer development (47), it will be interesting to investigate the phenotype induced by these mutations on the transcriptional and oncogenic function of TP53. Furthermore, as the characterisation of PTM modifications that enhance tumorigenecity of TP53 mutant protein has emerged in the last decade (43), it also raises the possibility that inhibition of T284 phosphorylation in the R282W mutant of TP53 contributes to the acquisition of its oncogenic function. By highlighting clusters of mutations that impact a common PTM site, our database provides hypotheses to experimentally investigate the role of this PTM in regulating the wildtype and the gain-of-function mutant TP53.

**Table 2.** PTM-associated cancer mutations affecting the phosphosite T284 in the tumor suppressor protein TP53. The table shows counts of somatic and germline disease mutations potentially affecting the phosphorylation site of Aurora kinase B in TP53.

| Phosphosite | Reference a.a. | Mutated a.a | Number of mutations |  | Impact on PTM |
| --- | --- | --- | --- | --- | --- |
|  |  |  | TCGA | ClinVar |  |
| T284 | R280 | T | 21 | 3 | distal |
|  |  | I |  | 3 | distal |
|  |  | K | 17 |  | distal |
|  |  | S | 3 |  | distal |
|  |  | G | 6 |  | distal |
|  | D281 | V | 4 | 2 | proximal |
|  |  | G |  | 2 | proximal |
|  |  | A | 2 |  | proximal |
|  |  | H | 4 |  | proximal |
|  |  | N | 3 |  | proximal |
|  |  | Y | 8 |  | proximal |
|  |  | E | 6 |  | proximal |
|  | R282 | W | 91 | 5 | network-rewiring |
|  |  | G | 4 | 5 | network-rewiring |
|  |  | Q | 1 | 2 | network-rewiring |
|  |  | L |  | 2 | network-rewiring |
|  | R283 | H | 1 | 2 | proximal |
|  |  | P | 9 |  | proximal |
|  |  | C |  | 3 | proximal |
|  |  | S |  | 3 | proximal |
|  | T184 | I |  | 1 | direct |
|  |  | P | 1 |  | direct |
|  |  | A |  |  | direct |
|  |  | E |  |  | direct |
|  | E285 | V | 4 | 2 | proximal |
|  |  | K | 25 |  | proximal |
|  | E286 | Q | 1 |  | network-rewiring |
|  |  | K | 17 | 1 | network-rewiring |
|  |  | A | 4 |  | proximal |
|  |  | G | 6 |  | proximal |
|  |  | V | 1 |  | proximal |
|  |  | Q |  | 1 | proximal |
| Total | - | - | 239 | 37 | - |

### Inherited and somatic PTM-associated mutations of BRCA2 and the DNA damage response pathway

Mutations in the tumor suppressor BRCA2 are linked to an elevated risk of breast and ovarian cancers as well as with Fanconi Anemia (FA), a rare chromosome instability syndrome characterized by aplastic anemia in children and susceptibility to leukemia and other types of cancer (48,49). Consistently, disease-associated SNVs of BRCA2 reported in the ClinVar database are associated with familial breast cancer and hereditary cancer-predisposing syndrome. At the molecular level, BRCA2 is centrally involved in DNA double strand break (DSB) repair by homologous recombination and other aspects of genome stability such as DNA replication, telomere homeostasis and cell cycle progression (50). To safeguard genomic stability at DSBs and stalled replication fork, BRCA2 relies on its ability to interact with RAD51, an interaction that is regulated by cell cycle-dependent kinases (CDKs) (51-54).

Using the ActiveDriverDB database, we studied PTM-associated SNVs in BRCA2 detected in the TCGA project and inherited disease mutations annotated in the ClinVar database. We found that a significant number of somatic and inherited cancer mutations coincide with phosphorylation sites (29 SNVs in ClinVar, FDR=10^−47^; 15 SNVs in TCGA, FDR=10^−6^)(**Figure 3**). One cluster of three phosphorylation sites (S3291, S3319, T3323) is located in the C-terminal of BRCA2, a region whose deletion is associated with increased radiation sensitivity and early onset of breast and ovarian cancers (55-57). The C-terminal domain of BRCA2 also contains nuclear import signals and a TR2 region (a.a. 3265-3330) that interact with nucleofilaments of RAD51 (52,54). Phosphorylation of TR2 by cyclin dependent kinases (CDKs) inhibits its ability to bind RAD51 nucleofilaments and is essential for entry into mitosis (51,52,54,58). Interestingly, the substitution of S3291A does not impact the DNA repair function of BRCA2 but it abolishes its ability to stabilize stalled replication forks (59). This observation suggests that this phosphorylation site plays an additional role during replication fork recovery. Substitutions that impact directly conserved phosphorylation sites (S3921, S3319, T3323) and CDK PTM consensus sites (P3292L/S, P3320H and P3324L) are associated with familial breast cancer and hereditary cancer-predisposing syndrome. Thus, we propose that mutations in this cluster of phosphosites abrogate the ability of BRCA2 to safeguard genomic stability either by inhibiting the interaction of BRCA2 with RAD51 nucleofilaments (S3291) or by a yet unknown mechanism. As the substitutions of S3319 and T3323 for glutamate do not abolish the interaction of TR2 with RAD51 in GST pull down assays (51), it is possible that these mutants interfere with the function of BRCA2 at stalled forks. Our analysis also highlights a substitution that potentially rewires phosphorylation events in the C-terminus of BRCA2. The mutation Q3321E creates a putative phosphorylation site for PLK1 (Polo-like kinase 1) that regulates multiple processes during mitosis. Disruption of BRCA2 during cytokinesis leads to genomic instability through a yet undefined mechanism (reviewed in (60)). During this process, PLK1 promotes the recruitment of BRCA2 to the midbody by phosphorylating its N-terminal region (S193) (61). As the functions of BRCA2 are differently regulated in a cell cycle dependent manner by CDKs and PLK1, aberrant phosphorylation of BRCA2 could impact genomic stability by rewiring its activities throughout different phases of the cell cycle. This example illustrates the integration of PTM information and genetic mutations to predict novel experimentally testable hypotheses of disease mechanisms.

**Figure 3.**
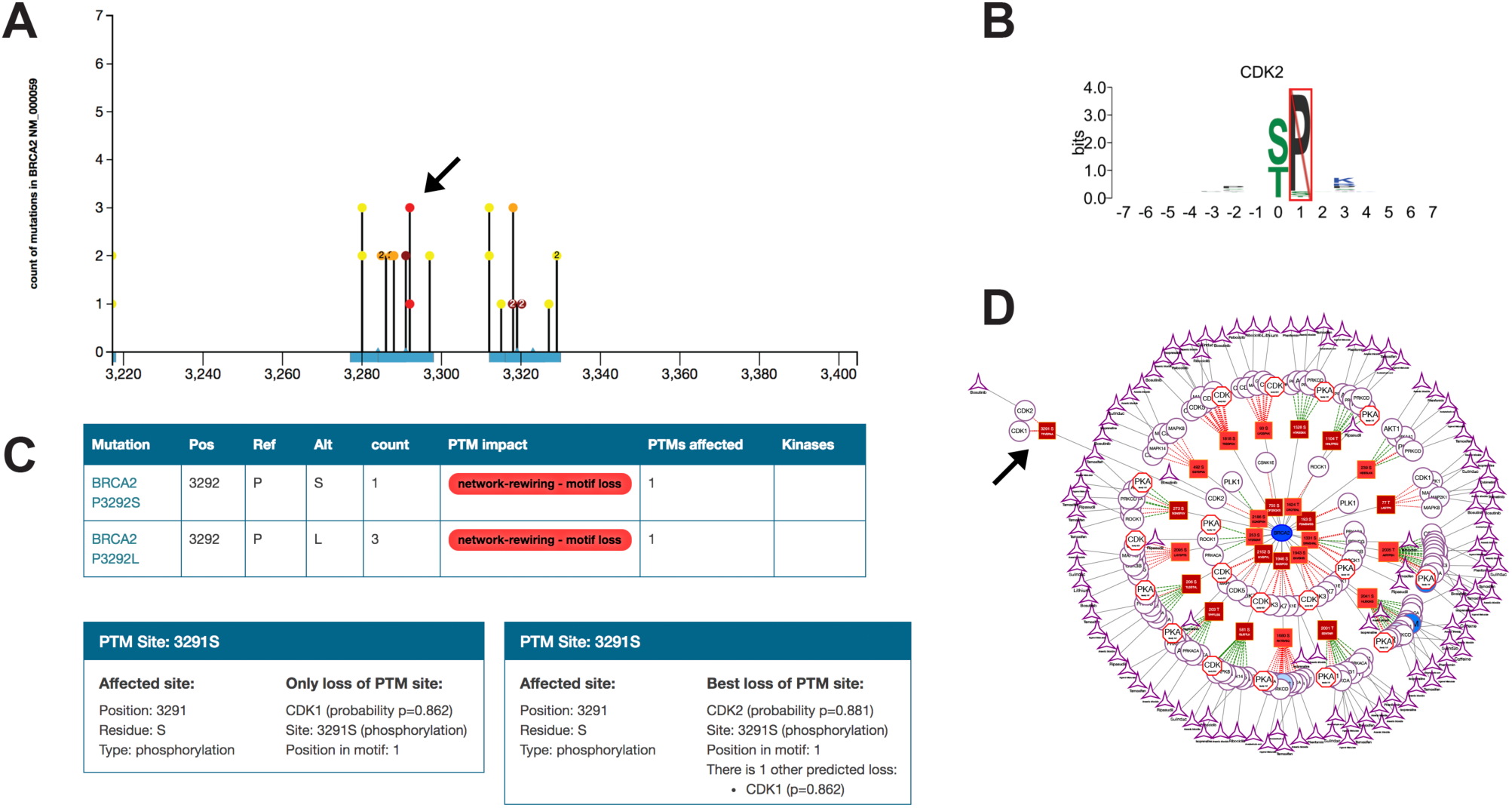
PTM-associated mutations in the BRCA protein involved in DNA repair and breast cancer. **(A)** Zoomed needleplot shows germline disease mutations from the ClinVar database located in three phosphorylation sites in the protein sequence at 3,200-3,400 residues. Only PTM-associated mutations are shown. **(B)** Mutations P3292L and P3292S are predicted to disrupt the sequence motif of the CDK2 kinase. **(C)** Table rows showing additional information on the two mutations. **(D)** The computationally predicted interaction network of BRCA2, its PTM sites, kinases predicted to interact with mutant BRCA2 according to the MIMP method, and drugs targeting these kinases. Arrows indicate the mutations of the residue P3292 in the protein sequence and interaction network.

### Network-rewiring mutations in the tumor suppressor VHL alter putative binding sites of CDK kinases

VHL is a tumor suppressor protein best known for its role in regulating cellular response to changes in tissue oxygen saturation (62). It encodes is the substrate recognition component of an E3 ubiquitin ligase complex that functions to constitutively down-regulate proto-oncogenic substrates such as protein kinase C, retinol binding protein 1, and hypoxia-inducible transcription factors (HIF) (62,63). VHL is frequently inactivated in cancer and up to 90% of all clear cell renal cell carcinomas (ccRCCs) harbour gene-silencing mutations (63). In particular, the mutation L169P along with the other p.157–172 subdomain mutations are considered hotspot mutations in ccRCCs (63) (**Figure 4**). Phosphorylation of VHL at S168 by NIMA Related Kinase 1 (NEK1) has been associated with VHL degradation and cilliary homeostasis (64). The mutation L169P is adjacent to the phosphosite S168 and another NEK1 phosphorylation site at Y175 and thus may impact the phoshposignalling of VHL.

**Figure 4.**
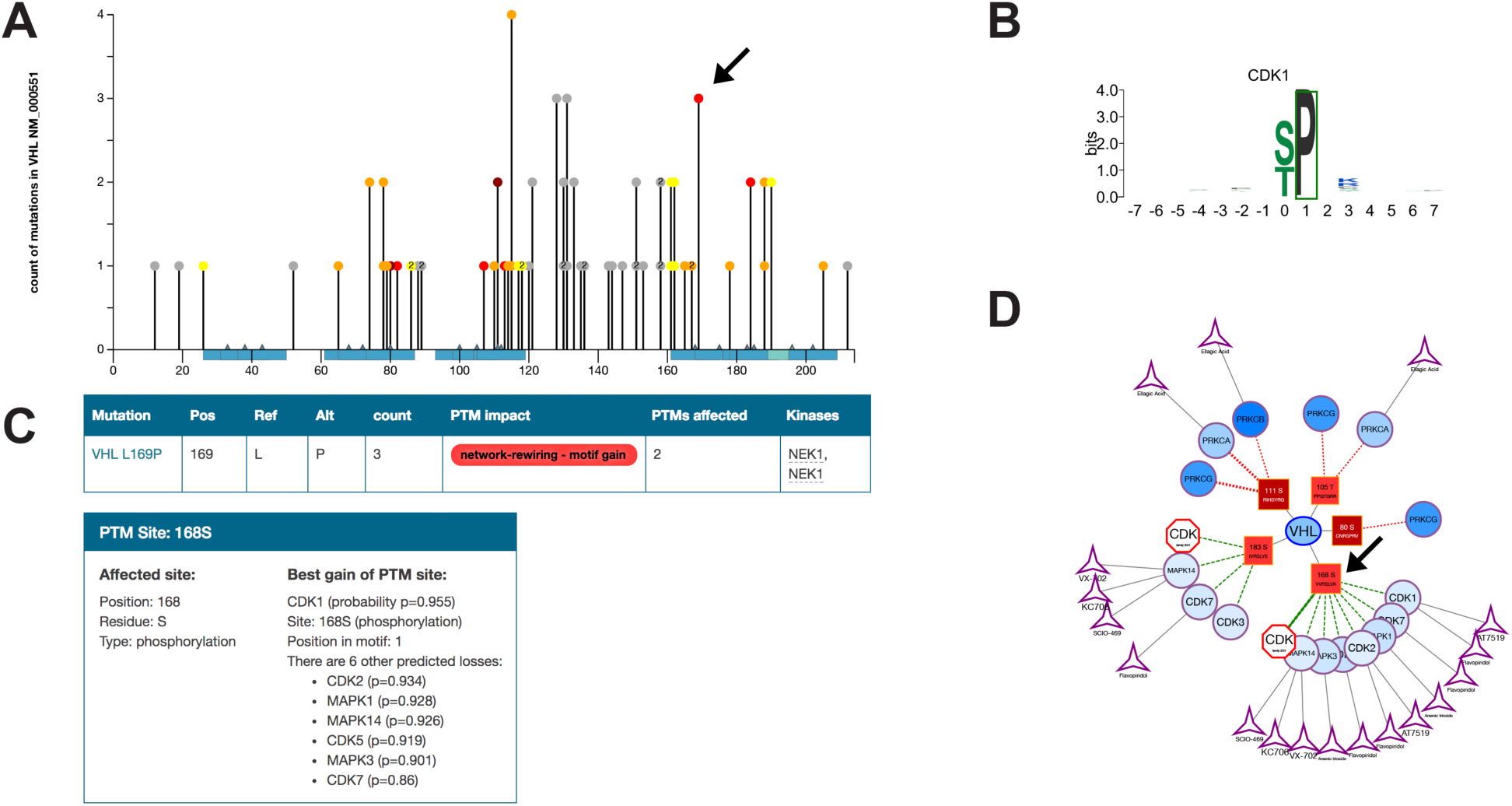
PTM-associated mutations in the tumor suppressor protein VHL. **(A)** Needleplot of mutations from the TCGA dataset with the mutation L169P located near two phosphorylation sites. **(B)** The mutations L196P is predicted to disrupt the sequence motif of the CDK1 kinase. **(C)** Table shows additional information on the mutation. **(D)** The computationally predicted interaction network of VHL, its PTM sites, kinases predicted to interact with mutant VHL according to the MIMP method, and drugs targeting these kinases. Arrows indicate the mutation L169P in the protein sequence and interaction network.

Given the mutation and kinase-network information in ActiveDriverDB, we propose that three L169P mutations observed in TCGA kidney cancers may cause gains of phosphorylation sites bound by the cyclin dependent kinase 1 (CDK1) or related kinases CDK2, CDK5, CDK7 and MAPK1, MAPK3 and MAPK14. While little is known about direct interactions between the oncogenic CDKs and VHL, VHL inactivation has been clinically linked to increased levels of CDK1 and CDK2 (65). CDK1 is also known to stabilize members of the HIF family, which are targets of VHL (66).

Investigating the role of the L169P mutation in the context of VHL phosphorylation and upstream kinases may reveal details about disease mechanisms. Since CDK1 is a drug targetable enzyme, assays with drugs such as alsterpaullone and alvocidib (33) may also be beneficial for research into targeted therapies.

## Discussion

ActiveDriverDB is a comprehensive human proteo-genomics resource that uses information on post-translational modifications of proteins to interpret disease mutations and inter-individual variation. While PTMs are important regulators of protein function with key roles in cellular signaling pathways, information about PTM sites is often not used for studying genetic variation and prioritizing candidate disease mutations. Our database aims to advance analysis of missense SNVs using PTM information and provides tools to users with different interests. Novice users of the database can start from example queries of well-annotated genes and browse lists of disease genes with significantly enriched PTM-associated SNVs. Basic and translational researchers can look up their favorite genes and pathways using various search options, and export tables and publication-quality figures stemming from their analysis. Geneticists can upload and interpret their candidate variants from DNA sequencing experiments and compare their findings with existing public datasets. Computational biologists and data scientists can use the ActiveDriverDB API to automatically analyze their variants of interest and download entire datasets for advanced studies.

We plan several important future developments of the database. Maintaining timely biomedical data resources is essential as new datasets accumulate rapidly and the use of outdated resources hampers scientific advances (67). Thus we aim to provide at least annual updates of our database to include recent large-scale genomics and proteomics studies and molecular interaction networks. In particular, advances in proteomics technologies enable large-scale characterization of other PTM types such as glycosylation (68) and SUMOylation (69) in human proteins and such emerging datasets will become available in future releases of the database. Datasets of additional species and genomes will be also considered, such as the most recent version of the human genome assembly (GRCh38) and model organisms such as mouse and Arabidopsis with abundant genome variation and proteomics data (14,70).

Interpreting inter-individual genetic variation will become an increasingly important challenge as we enter the era of personal genomics. Integration of proteomic and genomic information for deciphering the impact of variation on cellular systems and organism-level phenotypes is a powerful approach that will improve with the emergence of future datasets of increasing magnitude and complexity. We aim to provide an integrated database resource to the research community to interrogate these datasets and enable future discoveries.

## Acknowledgements

We would like to thank Andrea Sabo for help with web design. This work was supported by the Investigator Award to J.R. from the Ontario Institute for Cancer Research, the stipend to M.K. from the Google Summer of Code (GSoC) project, and Canadian Cancer Research Society (CSR) Operating Grant no. 21428 to A.F.T. and J.R. The results published here are in part based upon data generated by the TCGA Research Network: http://cancergenome.nih.gov/ as outlined in the TCGA publications guidelines http://cancergenome.nih.gov/publications/publicationguidelines. We are grateful to researchers and developers of databases PhosphoSitePlus, PhosphoELM, HPRD, ClinVar, DrugBank and others for providing high-quality and frequently maintained datasets.

